# SHARQnet - Sophisticated Harmonic Artifact Reduction in Quantitative Susceptibility Mapping using a Deep Convolutional Neural Network

**DOI:** 10.1101/522151

**Authors:** Steffen Bollmann, Matilde Holm Kristensen, Morten Skaarup Larsen, Mathias Vassard Olsen, Mads Jozwiak Pedersen, Lasse Riis Østergaard, Kieran O’Brien, Christian Langkammer, Amir Fazlollahi, Markus Barth

**Affiliations:** Centre for Advanced Imaging, The University of Queensland, Building 57 of University Dr, St Lucia QLD 4072, Brisbane, Australia; Department of Health Science and Technology, Aalborg University, Fredrik Bajers Vej 7, 9000, Aalborg, Denmark; Siemens Healthcare Pty Ltd, Brisbane, Australia; Department of Neurology, Medical University of Graz, Auenbruggerplatz 22, 8036, Graz, Austria; CSIRO Health and Biosecurity Flagship, The Australian eHealth Research Centre

**Author notes:** **Corresponding author:** Steffen Bollmann, Centre for Advanced Imaging, The University of Queensland. **Competing Interests statement**: Authors SB, MB, KO are co-inventors of a patent “Solving the ill-posed quantitative susceptibility mapping inverse problem using deep convolutional neural networks”, filed on 29th Dec 2017. KO is employed by Siemens Healthineers.

**Keywords:** Quantitative Susceptibility Mapping, Background field correction, Deep Learning

## Abstract

Quantitative susceptibility mapping (QSM) reveals pathological changes in widespread diseases such as Parkinson’s disease, Multiple Sclerosis, or hepatic iron overload. QSM requires multiple processing steps after the acquisition of magnetic resonance imaging (MRI) phase measurements such as unwrapping, background field removal and the solution of an ill-posed field-to-source-inversion. Current techniques utilize iterative optimization procedures to solve the inversion and background field correction, which are computationally expensive and lead to suboptimal or over-regularized solutions requiring a careful choice of parameters that make a clinical application of QSM challenging. We have previously demonstrated that a deep convolutional neural network can invert the magnetic dipole kernel with a very efficient feed forward multiplication not requiring iterative optimization or the choice of regularization parameters. In this work, we extended this approach to remove background fields in QSM. The prototype method, called SHARQnet, was trained on simulated background fields and tested on 3T and 7T brain datasets. We show that SHARQnet outperforms current background field removal procedures and generalizes to a wide range of input data without requiring any parameter adjustments. In summary, we demonstrate that the solution of ill-posed problems in QSM can be achieved by learning the underlying physics causing the artifacts and removing them in an efficient and reliable manner and thereby will help to bring QSM towards clinical applications.

## Introduction

Quantitative susceptibility mapping (QSM) is a post-processing technique that extracts magnetic susceptibility from the phase of magnetic resonance imaging (MRI) signal [1,2] and provides information about biological tissue properties, predominantly myelin [3], iron [4] and calcium [5]. QSM has been used to study normal aging [6], Huntington’s Disease [7], Multiple Sclerosis [8], Alzheimer’s Disease [9] and Parkinson’s Disease [10] and allows unambiguous visualization [11] and differentiation of micro-bleeds from microcalcifications [12].

Obtaining a quantitative susceptibility map requires an MRI sequence where the signal phase is sensitive to local magnetic field changes, such as a gradient-recalled-echo sequence [13] or gradient-echo-based echo-planar imaging [14,15]. This raw signal phase is first unwrapped, and the background field, magnetic field changes from regions outside the object of interest, have to be removed before the measured field perturbation can be related to the underlying tissue magnetic susceptibility distribution by solving an ill-posed inverse problem [1].

The background field in QSM is caused by magnetic field changes outside the object of interest, such as susceptibility gradients due to tissue-air interfaces [16] or B_0_ inhomogeneities due to imperfect shimming. These external fields are often orders of magnitude stronger and overlap with local tissue field changes. A variety of methods have been proposed to remove background fields in QSM by exploiting the underlying physical principle that the background phase either satisfies the Laplace equation inside the object of interest or that the background phase is caused by sources outside the object of interest. This means that the internal tissue-related fields can be modeled as non-harmonic components, whereas background fields can be modeled as harmonic components of the total field perturbation. A recent review [17] classified background field corrections based on their assumptions in 1) methods assuming no sources close to boundaries, 2) methods assuming no harmonic internal and boundary fields, and 3) methods that do not employ an explicit boundary assumption, but minimize an objective function based on a norm. One example for satisfying the Laplace equation inside the object of interest, assuming no sources close to boundaries, is sophisticated harmonic artifact reduction for phase data (SHARP) [16]. It solves Poisson’s equation utilizing the spherical mean value theorem [18] and requires the definition of a spherical kernel radius and a regularization parameter. The SHARP method was later extended to V-SHARP, utilizing spheres with multiple radii that decrease the size of the kernel towards the brain boundary to reduce artifacts at the edges and thereby extend the usable region of interest [19]. The most recent SHARP scheme utilizes a high-pass filter to define the regularization parameters more robustly [20], allowing the use of the same regularization values for different imaging parameters.

Another method, assuming no harmonic internal and boundary fields, removes the background field by solving the Laplacian boundary value problem (LBV) [21]. Under simple boundary condition assumptions, this method removes the background field while retaining data near the boundary. The simple boundary condition assumptions work in most cases, but can be problematic when the local field is very high near the boundary (e.g. veins close to the brain surface).

One example in the third group is regularization enabled SHARP (RESHARP) [22], which introduces a Tikhonov regularization at the deconvolution stage. Methods that are based on physical properties of dipole sources outside the object of interest, such as projection onto dipole fields (PDF) [23,24], also fall in this category because they fit a distribution of external sources to the total field by projecting the field inside the object onto the subspace spanned by all background sources.

In summary, the currently available background field corrections perform well given that the regularization parameters are carefully adjusted to a given dataset and the assumptions of the boundary conditions are not violated. Most methods face the limitation of a loss of information at the boundaries, which can be partly mitigated by using varying kernel sizes [25] or by extending field coverage exploiting the harmonic field properties [26]. A fundamental problem of most available methods is that they require the definition of a mask, separating the object of interest from the background. However, this mask generation is non-trivial (especially in the abdomen or heart) and leads to either the loss of areas close to boundary regions or residual artifacts due to an incorrect boundary definition. The need for carefully choosing regularization parameters and defining a region of interest currently limit the wide clinical application of QSM.

Deep convolutional neural networks have recently been shown to enable an efficient solution of the ill-posed inversion problem [27,28] without requiring computationally expensive iterative optimization procedures or the explicit choice of regularization parameters. We extended the previously introduced framework DeepQSM [27] to remove background fields by learning to predict realistic background fields from simulated data. In this manuscript, we describe the simulation of background fields used for training and compare the predictions of our prototype method SHARQnet with established background field removal procedures.

## Methods

### Training data

For training SHARQnet, we simulated 1000 realistic background fields overlaid on top of a synthetic brain simulation provided with the MEDI toolkit [29]. This was achieved by randomly placing 2 or 3 ellipsoids of varying sizes outside the brain phantom and simulating the fields originating from these ellipsoids by convolving them with the unit dipole response. The external sources varied in susceptibility and shape. Then, a brain mask was applied to the brain phantom with the overlaid simulated background fields, and 100 different crops (32×32×32) were randomly extracted (allowing random overlap between patches) to yield the input data for SHARQnet (Figure 1).

**Figure 1.**
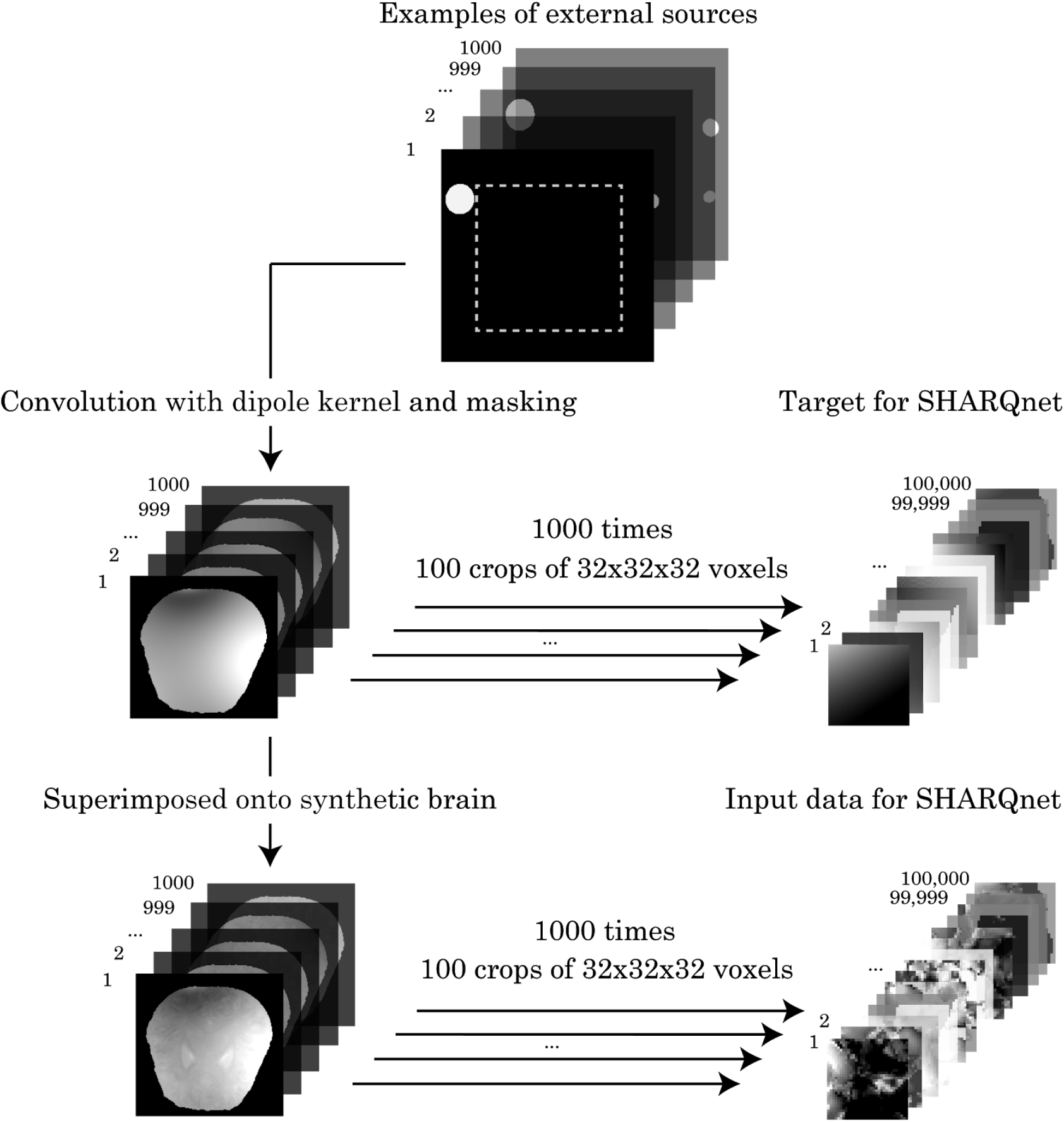
Illustration of the training data generation for SHARQnet. 1 000 different source configurations were simulated, convolved with the dipole kernel and superimposed onto a synthetic brain model. 100 different examples of size 32×32×32 were randomly cropped from each brain resulting in 100 000 examples used for training.

The training target was the synthetic background field, which was achieved by subtracting the output of the network from the input and computing the mean squared error with respect to the brain phantom without background fields (for training details see Figure 2).

**Figure 2.**
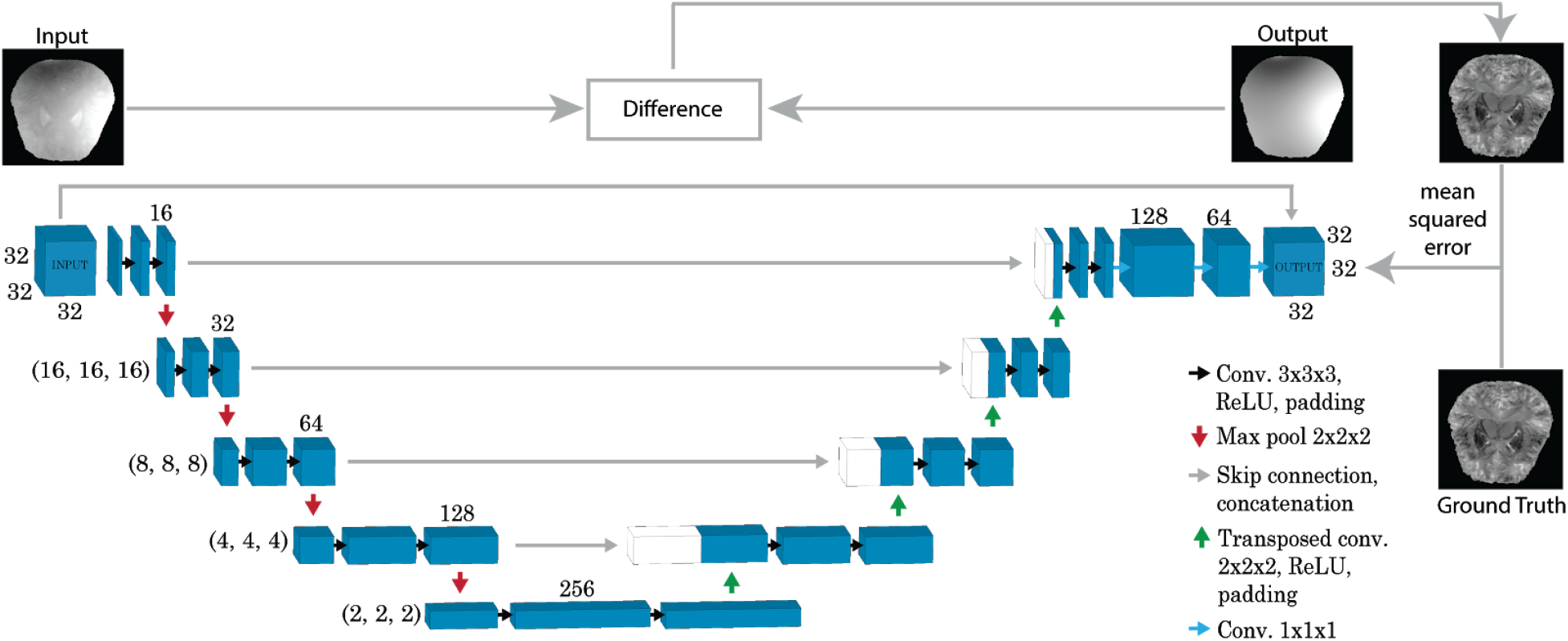
Network architecture of SHARQnet. The input image in this example has a dimension of 32×32×32 and is processed by two convolutional and one max pooling layer at each level before the image is upsampled via transposed convolutional layers followed by two convolutional layers. Finally, the output is processed via two adaptation layers with a kernel size of 1. The output of the network is then subtracted from the input, and the mean squared error is computed between this difference image and the ground truth image. The error is used for adjusting the training weights during back propagation.

### Architecture and training

The fully convolutional neural network is based on DeepQSM [27], which is a modified version of an U-Net architecture [30]. Due to memory constraints, we reduced the amount of feature maps compared to the original U-Net (Figure 2). In addition to the DeepQSM architecture, we utilized two adaptation layers with 128 and 64 filters respectively and a kernel size of 1. The network expects 3D background field superimposed images as input and produces an equally sized predicted background field as output.

SHARQnet was trained on 100 000 synthetic background examples with a mini-batch size of 120 in a total training time of 24 hours on an NVIDIA Tesla K40 Accelerator Unit. To optimize the weights of SHARQnet during training, the ‘ADAM’ optimizer [31] was used with default parameters. To avoid overfitting during training, the regularization technique ‘dropout’ was used [32] to randomly turn off neurons during training with a dropout rate of 15%. This dropout rate was chosen, because commonly used higher rates of 50% increased the training time considerably and we were able to simulate large amounts of data and by this avoiding overfitting more efficiently. We always started the training from scratch and did not use pre-trained networks. The network expects input data that are independent of echo time and field strength and this normalization step has to be applied beforehand. Due to the quantitative nature of the QSM problem, it is not necessary to apply sophisticated normalization techniques as in the case of image segmentation problems utilizing T_1_ or T_2_ weighted data [33].

SHARQnet was implemented using Python 3.6 [34], Tensorflow and Tensorboard v1.6 [35], Keras 2.2.4 [36], SciPy 1.1.0 [37], NumPy 1.15.0 [38], scikit-image 0.13.1 [39] and Nibabel 2.3.0 [40]. Matplotlib 2.1.2 [41] was used for visualizations and creating figures in this manuscript. The source code of SHARQnet is available from the authors upon request. Training was performed on the National Computational Infrastructure cluster Raijin.

Three experiments were performed to evaluate SHARQnet’s ability to remove background fields. As a comparison, we used three established background field correction methods and we reconstructed the same datasets using SHARP [16], RESHARP [22] and V-SHARP [19].

### Performance evaluation 1: Synthetic data

The first experiment aimed to test SHARQnet’s performance on synthetic background fields similar to the training data. The outputs of SHARQnet were compared to the ground truth through the use of difference maps and numerical error measures, such as the normalized root-mean-squared error (RMSE), the structural similarity index (SSIM [42]), and the high-frequency error norm (HFEN [43]).

### Performance evaluation 2: QSM Reconstruction Challenge data

In the second experiment, the goal was to remove the background field from realistic single-orientation phase data. SHARQnet had never been introduced to real background fields during training, and therefore this experiment tests if the simulated background fields characterize real background fields well enough to enable a correct removal. The single transversal-orientated unwrapped phase image from the 2016 QSM reconstruction challenge was used. We chose this publicly available dataset because it resembles a typical clinical dataset and it enables everyone a comparison to their own in-house algorithms. The data set serves as a common reference for current and future algorithms and was acquired *in vivo* from a healthy 30-year-old female, using a 3D gradient-echo sequence at 3 T with 1.06 mm isotropic resolution, an echo time of 25 ms, and a repetition time of 35 ms [44]. The run time of SHARQnet is dependent on the input matrix size and is on average 42s on an i7-4790 CPU for this 160×160×160 dataset. To demonstrate that the background field corrected maps can be used in a QSM pipeline, we applied the DeepQSM [27] dipole inversion algorithm to the output of every background field correction algorithm.

### Performance evaluation 3: High-resolution 7T *in vivo* data with and without brain extraction

The third experiment tests if the background field can be removed from a dataset acquired at ultra-high field at a variety of echo times. In addition, we tested if SHARQnet can produce background field corrections without brain masking. For this, we obtained written informed consent from one participant (27 years, male) prior to *in vivo* scanning as approved by the local human ethics committee on a 7T whole-body research scanner (Siemens Healthcare, Erlangen, Germany), with maximum gradient strength of 70 mT/m and a slew rate of 200 T/m/s using the 32-channel Rx head coil (Nova Medical, Wilmington, MA, USA). Third-order shimming was employed to improve the B_0_-field homogeneity.

We acquired a multiple-echo-time gradient-recalled-echo (GRE) 3D whole-brain dataset: TR = 25 ms, TE = 4.4, 7.25, 10.2, 13.25, 16.4, 19.65, 23 ms, flip angle = 13°, FOV = 210×181.5×120 mm^3^, matrix = 280×242×160 (0.75 mm isotropic voxels), parallel imaging (GRAPPA, acceleration factor = 2, 24 auto-calibration lines), monopolar readout gradient, symmetric echo, 1116 Hz/Pixel, first echo flow compensated, TA = 7.9 min.

To enable optimal coil combination using COMPOSER [45], we acquired reference data using the prototype PETRA ultra-short-TE sequence [46]: TR = 1.99 ms, TE = 0.07 ms, flip angle = 2°, FOV = 288×288×288 mm^3^, matrix = 288×288×288 (1 mm isotropic voxels), 847 Hz/Pixel, and TA = 2 min.

The data was cropped to 224×272×160 pixels, and a brain mask was generated using Oxford FMRIB Software Library (FSL) Brain Extraction Tool (BET) [47] with a fractional intensity threshold of 0.4 per individual echo. Unwrapping was performed using STI Suite’s Laplacian unwrapping algorithm [48]. All echoes were individually processed with and without brain masking.

## Results

### Performance evaluation 1: Synthetic data

Figure 3 shows the results of SHARQnet, SHARP, RESHARP and V-SHARP on simulated background fields overlaid onto a simulated brain. Although all methods are capable of removing the background field, the comparison to the ground truth reveals the consequences of the underlying assumptions of each method. SHARP produces a relatively flat image and suppresses a wide range of spatial frequencies including fields generated from structures of interest inside the brain. Not unexpected given their shared concept, RESHARP and V-SHARP show residuals with lower spatial frequencies. However, all three methods show errors at the brain boundary, which can be seen in the top part of the brain in the coronal slices. SHARQnet in comparison shows substantially smaller errors, manifested predominantly in high spatial-frequency components. The numerical error measures (Table 1) affirm the visual comparisons and indicate that SHARQnet performs best.

**Figure 3.**
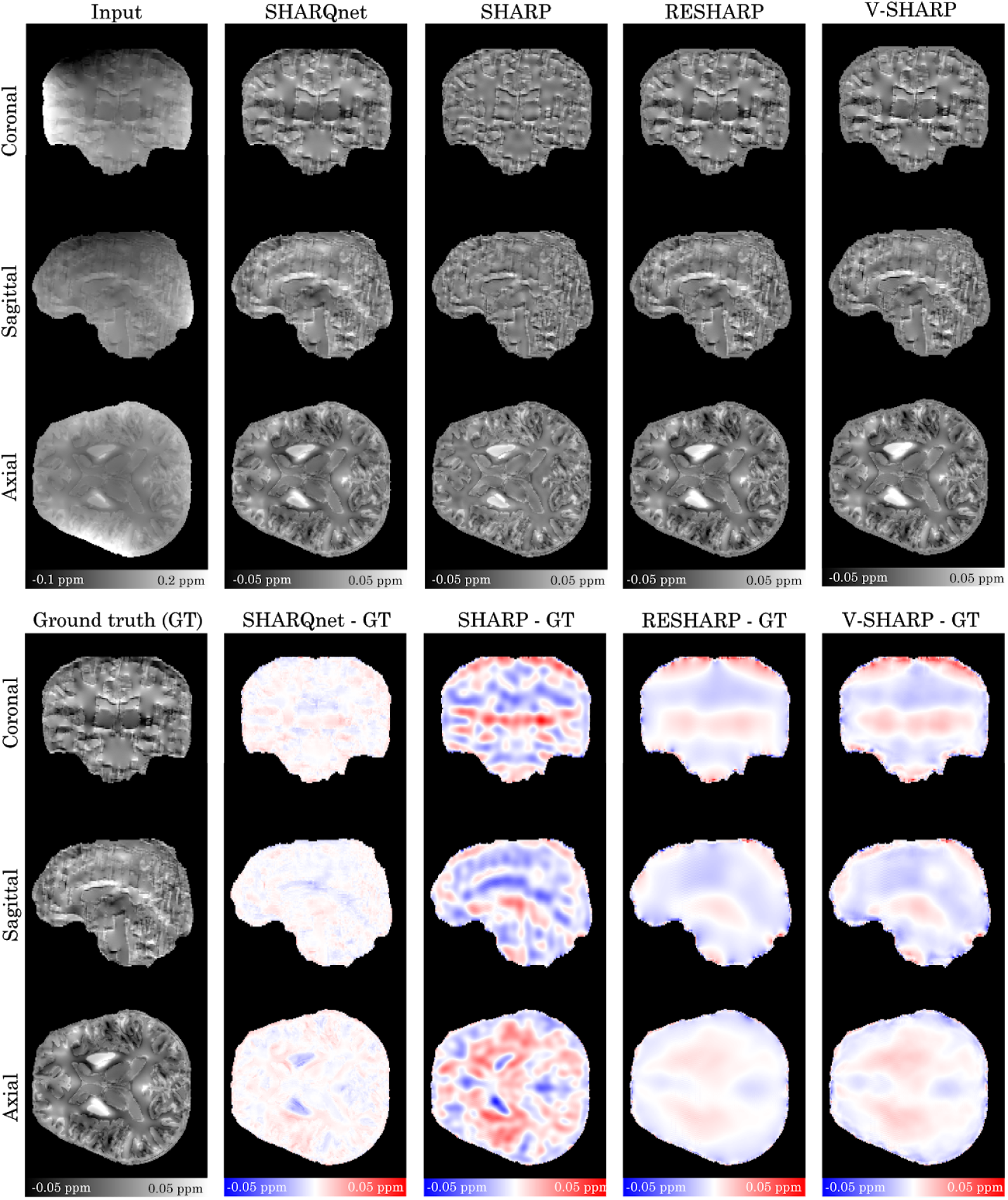
Results of SHARQnet, SHARP, RESHARP and V-SHARP background field removal on synthetic data (top row) as well as the difference to the ground truth (bottom row). The input is a simulated background field superimposed onto the simulated brain phantom. Compared to the SHARP-based methods, SHARQnet delivers substantially lower residual background fields.

**TABLE 1.** Error metrics of outputs from SHARQnet, SHARP, RESHARP and V-SHARP for background field removal on a synthetic brain with simulated background field: root mean square error (RMSE), high-frequency error norm (HFEN) and structural similarity index (SSIM) enable the examination of an overall error, deviation in high-frequency components, and visual similarity respectively. SHARQnet performs better than SHARP-based methods in all metrics used.

| Method | RMSE | HFEN | SSIM |
| --- | --- | --- | --- |
| SHARQnet | 19.55 | 17.01 | 0.9875 |
| SHARP | 70.37 | 51.35 | 0.8353 |
| RESHARP | 41.97 | 29.60 | 0.8811 |
| V-SHARP | 47.01 | 34.37 | 0.8735 |

### Performance evaluation 2: QSM Reconstruction Challenge data

The results of SHARQnet, SHARP, RESHARP and V-SHARP on the 2016 QSM Reconstruction Challenge data are shown in Figure 4. The yellow arrow highlights a region where SHARQnet and SHARP remove the background field visually correctly, but RESHARP and V-SHARP show a residual field. The red arrow highlights an area where SHARP removes the background field, whereas SHARQnet maintains more anatomical contrast from basal ganglia structures. It can also be seen that SHARQnet delivers a lower susceptibility for the superior sagittal sinus and straight sinus compared to the other methods. The QSM reconstruction of the background field corrected data is shown as well and it can be observed that SHARQnet delivers a background field correction that allows the computation of a susceptibility map of high fidelity.

**Figure 4.**
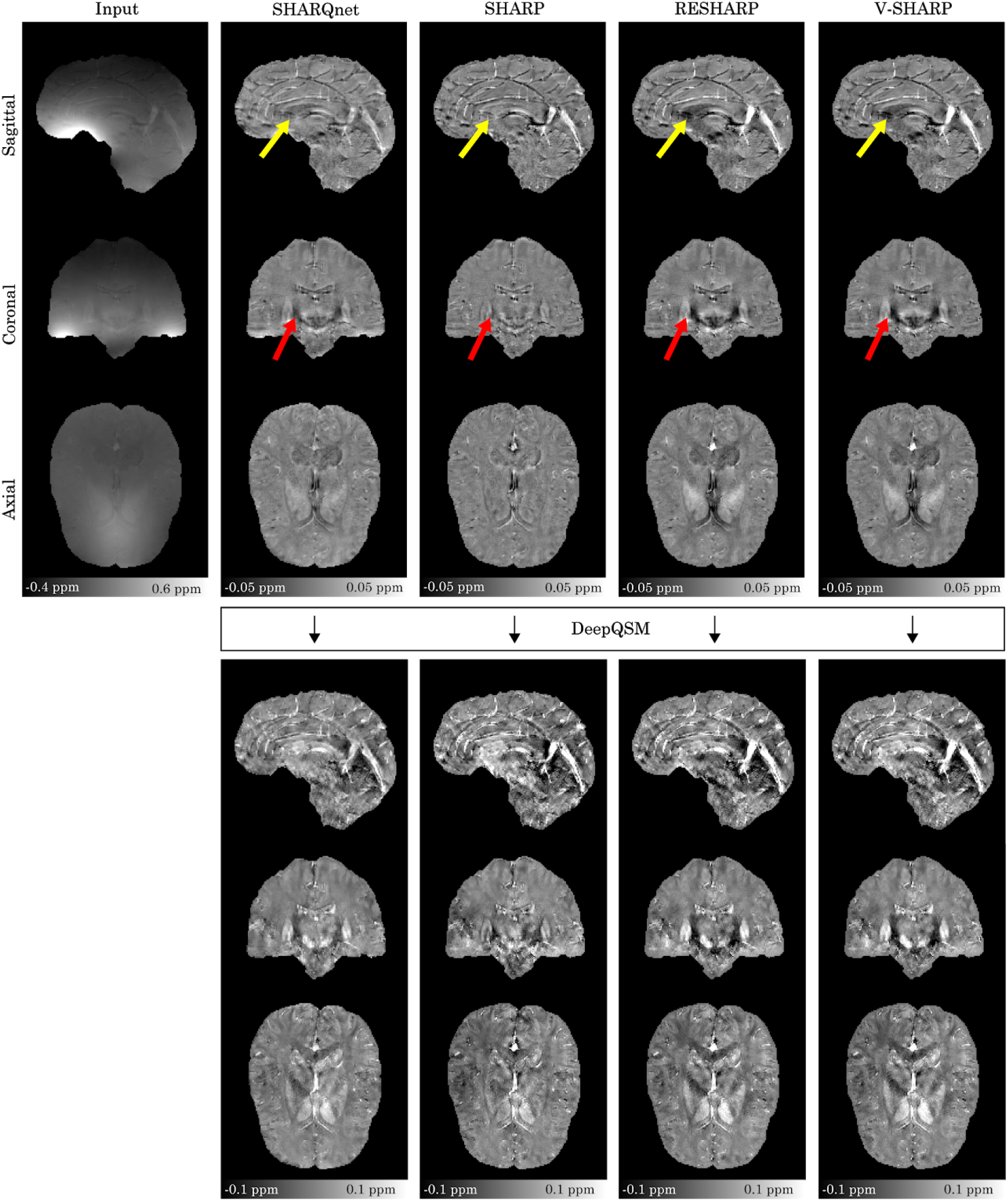
Results of the background field removal using SHARQnet, SHARP, RESHARP and V-SHARP on the 2016 QSM Reconstruction Challenge data. The input is the unwrapped phase from the challenge. The yellow arrow highlights a region where SHARQnet and SHARP remove the background field, but RESHARP and V-SHARP show a residual field. The red arrow points to a basal ganglia area where the background field removal by SHARP might lead to an underestimation of susceptibility, whereas SHARQnet, RESHARP and V-SHARP maintain anatomical contrast of those iron-rich structures. The second panel shows the result of the DeepQSM dipole inversion when the respective background field corrected images are utilized as input.

### Performance evaluation 3: High-resolution 7T *in vivo* data with and without brain extraction

Figure 5 and 6 show SHARQnet’s results on a 7T multi-echo dataset. Figure 5 demonstrates that SHARQnet is capable of removing the background field from an ultra-high field dataset across multiple echo-times. Figure 6 shows that SHARQnet even works without a brain mask defining the object of interest and delivers realistic background field corrections.

**Figure 5.**
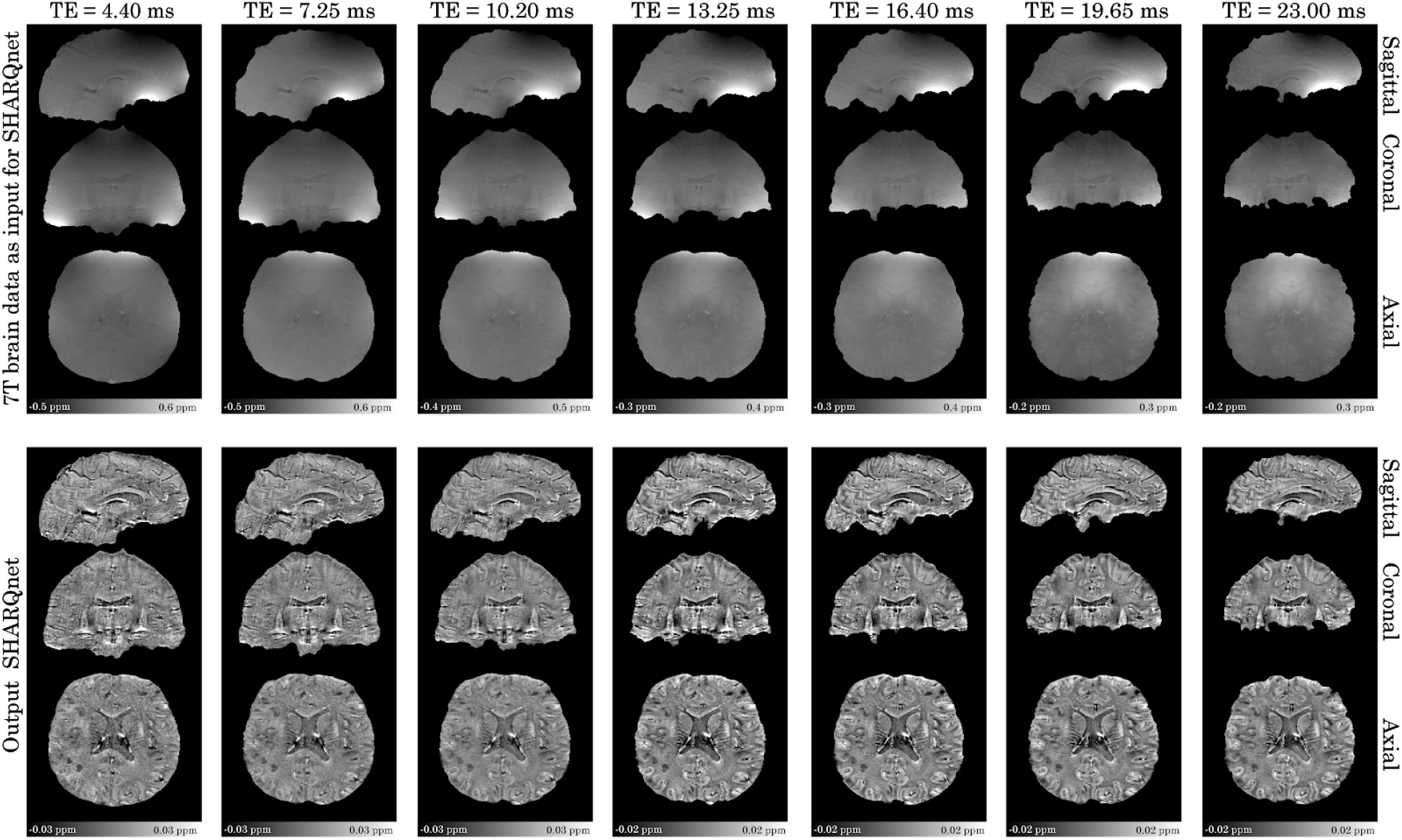
Results of SHARQnet on a 7T multi-echo dataset. The individual echoes have been masked with an echo-time-specific mask. SHARQnet is able to remove the background field in this dataset for every echo-time.

**Figure 6.**
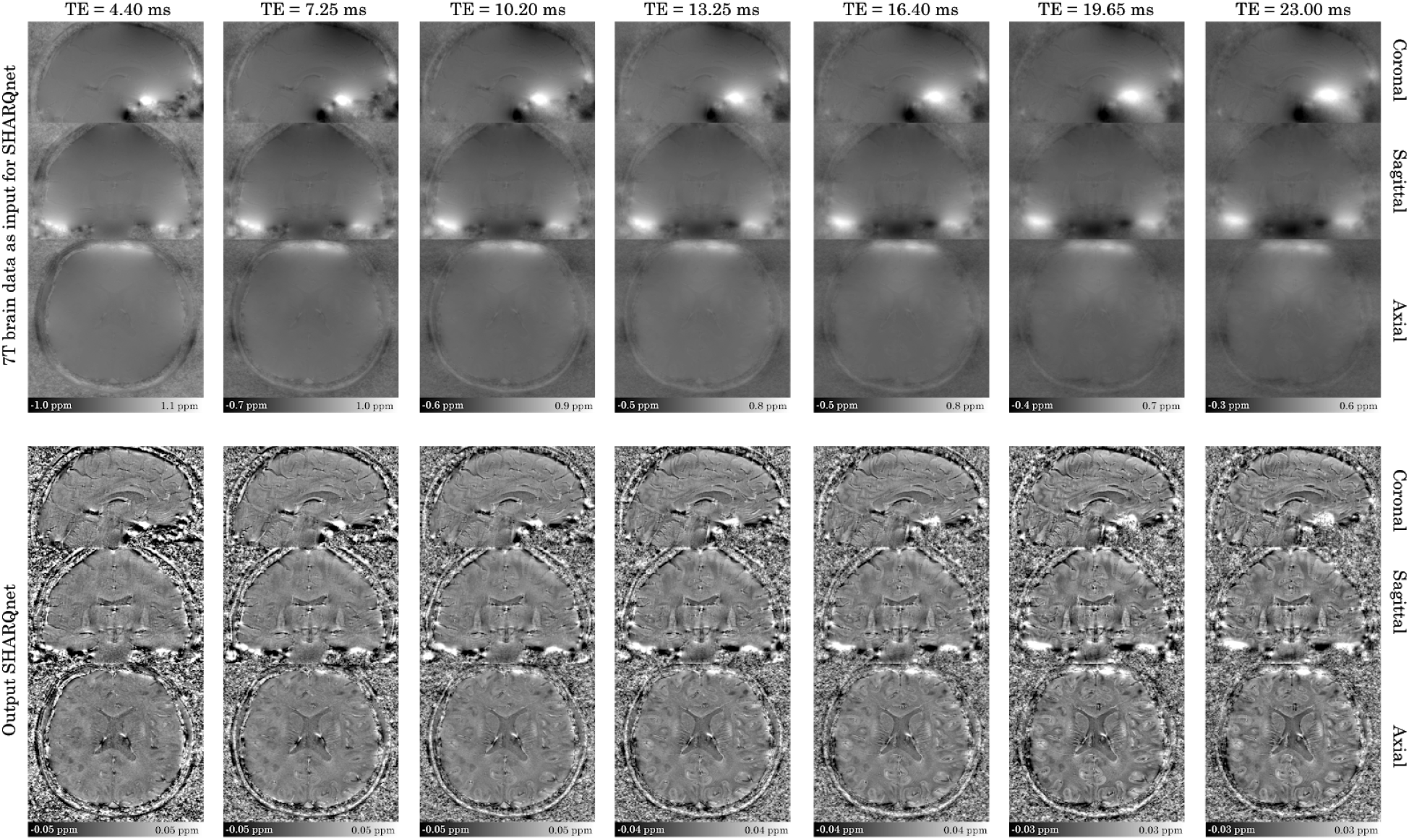
Results of SHARQnet on a 7T multi-echo dataset, without brain masking being applied. It can be seen that SHARQnet is able to remove the background field even without a mask defining the object of interest.

## Discussion

In this work, we introduced SHARQnet, a deep convolutional neural network that is capable of removing background fields in QSM. We compared SHARQnet with three commonly used algorithms SHARP, RESHARP and V-SHARP and found that it provides accurate background field corrections across all tested applications.

When testing SHARQnet on synthetic background fields we observed that it yields lower numerical error metrics than the state-of-the-art methods. The difference maps (Figure 3) facilitate interpreting the lower error metrics and indicate that SHARQnet only shows residual errors close to high susceptibility structures (e.g. the basal ganglia). Importantly, SHARQnet works well close to the brain boundary, where it shows substantially smaller errors in comparison to SHARP, V-SHARP and RESHARP.

We also tested SHARQnet on real-world data where no ground truth is available and found that SHARQnet produces artifact-free results that appear visually correct. In this test case, it is more difficult to evaluate the performance because there is no ground truth. However, when combining with the insights gained from the synthetic data, we found that SHARP might regionally reduce the anatomical contrast too excessively, potentially resulting in reduced magnetic susceptibility estimates in e.g. iron-rich basal ganglia structures. Additionally, the output of SHARQnet differs from the other methods in the large vein depicted in Figure 4, and it is likely that SHARQnet underestimates susceptibility in this structure. We speculate that our training dataset might not have contained enough structures with high susceptibility values and that the network therefore partly removes the field associated with veins. Future implementations should investigate this in more detail.

We also demonstrated that SHARQnet generalizes to *in vivo* data acquired at ultra-high field at varying echo times. Furthermore, we show that SHARQnet does not necessarily need a mask defining the object of interest like the competing algorithms. If a further augmentation of the source simulation, e.g. by adding sources close to the temporal lobe, can help to eliminate the residual fields in the temporal lobes (e.g depicted in Figure 6), it might even be possible to omit the error-prone brain masking step from the QSM pipeline. This would have huge advantages in clinical applications as has already been shown in QSM algorithms that can invert the total field [49,50] and do not require brain masking.

The presented work is a proof-of-concept that has huge potential to be improved further. First, the simulation of the external background fields is relatively simple and could be done in a more anatomically realistic way. The simple source geometry currently used, could explain the residual fields visible in the temporal lobes. For applications outside the brain, such as QSM in the abdomen [51], it needs to be tested if the current simulation approach already generalizes to other organs of interest or if organ-specific simulations should be utilized. More room for improvement can be found in the network architecture and hyperparameters, because we currently have not optimized these extensively. Preliminary tests (not shown in this manuscript) indicate that smaller architectures are capable of learning the background field problem as well. In conclusion, we present a novel background field removal technique in QSM based on a deep convolutional neural network. We show that the presented algorithm delivers numerically and visually better results than state-of-the-art methods, but does not require manual parameter tuning. Additionally, preliminary results indicate that our proposed method might not necessarily require brain or organ masking, rendering it especially interesting for clinical applications outside the brain.

## Acknowledgements

The authors acknowledge the facilities and scientific and technical assistance of the National Imaging Facility, a National Collaborative Research Infrastructure Strategy (NCRIS) capability, at the Centre for Advanced Imaging, The University of Queensland. MHK, MSL, MVO and MJP acknowledge financial support from the following private organisations: the Obel Family Foundation, the Knud Højgaard Foundation, the Augustinus Foundation, the Oticon Foundation, Otto Mønsted’s Foundation, the Henry and Mary Skov’s Foundation, Viet-Jacobsen Foundation, Torben and Alice Frimodt’s Foundation, Aalborg Stiftstidende Foundation, Dansk Tennis Foundation, Marie and M.B. Richters Foundation, Julie Damms Foundation, the Roblon Foundation, Vanggaard Foundation, William and Hugo Evers Foundation, the Frimodt-Heineke Foundation, Statsautoriseret Revisor Oluf Christian Olsen and Hustru Julie Rasmine Olsens Foundation, and Reinholdt W. Jorck and Hustrus Foundation. CL was supported by the Austrian Science Fund (FWF grant numbers: KLI523 and P30134). MB acknowledges funding from Australian Research Council Future Fellowship grant FT140100865. This research was undertaken with the assistance of resources and services from the Queensland Cyber Infrastructure Foundation (QCIF) and the National Computational Infrastructure (NCI), which is supported by the Australian Government. We would like to thank the reviewers for the constructive and very helpful feedback.

## References

[1] Deistung A, Schweser F, Reichenbach JR. Overview of quantitative susceptibility mapping: Overview of Quantitative Susceptibility Mapping. NMR Biomed 2017;30:e3569. doi:10.1002/nbm.3569.

[2] Schweser F, Deistung A, Reichenbach JR. Foundations of MRI phase imaging and processing for Quantitative Susceptibility Mapping (QSM). Z Für Med Phys 2015. doi:10.1016/j.zemedi.2015.10.002.

[3] Wharton S, Bowtell R. Effects of white matter microstructure on phase and susceptibility maps. Magn Reson Med 2015;73:1258–69. doi:10.1002/mrm.25189.

[4] Ropele S, Langkammer C. Iron quantification with susceptibility: Iron quantification with susceptibility. NMR Biomed 2016. doi:10.1002/nbm.3534.

[5] Buch S, Liu S, Ye Y, Cheng Y-CN, Neelavalli J, Haacke EM. Susceptibility mapping of air, bone, and calcium in the head. Magn Reson Med 2015;73:2185–94. doi:10.1002/mrm.25350.

[6] Acosta-Cabronero J, Betts MJ, Cardenas-Blanco A, Yang S, Nestor PJ. In Vivo MRI Mapping of Brain Iron Deposition across the Adult Lifespan. J Neurosci 2016;36:364–74. doi:10.1523/JNEUR0SCI.1907-15.2016.

[7] Bergen JMG van, Hua J, Unschuld PG, Lim I a. L, Jones CK, Margolis RL, et al. Quantitative Susceptibility Mapping Suggests Altered Brain Iron in Premanifest Huntington Disease. Am J Neuroradiol 2015. doi:10.3174/ajnr.A4617.

[8] Barkhof F, Thomas DL. Mapping Deep Gray Matter Iron in Multiple Sclerosis by Using Quantitative Magnetic Susceptibility. Radiology 2018:181274. doi:10.1148/radiol.2018181274.

[9] Acosta-Cabronero J, Williams GB, Cardenas-Blanco A, Arnold RJ, Lupson V, Nestor PJ. In Vivo Quantitative Susceptibility Mapping (QSM) in Alzheimer’s Disease. PLoS ONE 2013;8:e81093. doi:10.1371/journal.pone.0081093.

[10] Langkammer C, Pirpamer L, Seiler S, Deistung A, Schweser F, Franthal S, et al. Quantitative Susceptibility Mapping in Parkinson’s Disease. PLOS ONE 2016;11:e0162460. doi:10.1371/journal.pone.0162460.

[11] Liu T, Surapaneni K, Lou M, Cheng L, Spincemaille P, Wang Y. Cerebral microbleeds: burden assessment by using quantitative susceptibility mapping. Radiology 2012;262:269–78. doi:10.1148/radiol.11110251.

[12] Schweser F, Deistung A, Lehr BW, Reichenbach JR. Differentiation between diamagnetic and paramagnetic cerebral lesions based on magnetic susceptibility mapping. Med Phys 2010;37:5165–78. doi:10.1118/1.3481505.

[13] Reichenbach JR, Venkatesan R, Yablonskiy DA, Thompson MR, Lai S, Haacke EM. Theory and application of static field inhomogeneity effects in gradient-echo imaging. J Magn Reson Imaging 1997;7:266–79. doi:10.1002/jmri.1880070203.

[14] Langkammer C, Bredies K, Poser BA, Barth M, Reishofer G, Fan AP, et al. Fast quantitative susceptibility mapping using 3D EPI and total generalized variation. NeuroImage 2015;111:622–30. doi:10.1016/j.neuroimage.2015.02.041.

[15] Sun H, Wilman AH. Quantitative susceptibility mapping using single-shot echo-planar imaging. Magn Reson Med 2015;73:1932–8. doi:10.1002/mrm.25316.

[16] Schweser F, Deistung A, Lehr BW, Reichenbach JR. Quantitative imaging of intrinsic magnetic tissue properties using MRI signal phase: An approach to in vivo brain iron metabolism? NeuroImage 2011;54:2789–807. doi:10.1016/j.neuroimage.2010.10.070.

[17] Schweser F, Robinson SD, de Rochefort L, Li W, Bredies K. An illustrated comparison of processing methods for phase MRI and QSM: removal of background field contributions from sources outside the region of interest. NMR Biomed 2016:n/a–n/a. doi:10.1002/nbm.3604.

[18] Li L, Leigh JS. High-precision mapping of the magnetic field utilizing the harmonic function mean value property. J Magn Reson San Diego Calif 1997 2001;148:442–8. doi:10.1006/jmre.2000.2267.

[19] Li W, Wu B, Liu C. Quantitative susceptibility mapping of human brain reflects spatial variation in tissue composition. NeuroImage 2011;55:1645–56. doi:10.1016/j.neuroimage.2010.11.088.

[20] Özbay PS, Deistung A, Feng X, Nanz D, Reichenbach JR, Schweser F. A comprehensive numerical analysis of background phase correction with V-SHARP. NMR Biomed 2016. doi:10.1002/nbm.3550.

[21] Zhou D, Liu T, Spincemaille P, Wang Y. Background field removal by solving the Laplacian boundary value problem: BACKGROUND FIELD REMOVAL BY SOLVING LAPLACIAN BOUNDARY VALUE PROBLEM. NMR Biomed 2014;27:312–9. doi:10.1002/nbm.3064.

[22] Sun H, Wilman AH. Background field removal using spherical mean value filtering and Tikhonov regularization. Magn Reson Med 2014;71:1151–7. doi:10.1002/mrm.24765.

[23] de Rochefort L, Liu T, Kressler B, Liu J, Spincemaille P, Lebon V, et al. Quantitative susceptibility map reconstruction from MR phase data using bayesian regularization: Validation and application to brain imaging. Magn Reson Med 2010;63:194–206. doi:10.1002/mrm.22187.

[24] Liu T, Khalidov I, de Rochefort L, Spincemaille P, Liu J, Tsiouris AJ, et al. A novel background field removal method for MRI using projection onto dipole fields (PDF). NMR Biomed 2011;24:1129–36. doi:10.1002/nbm.1670.

[25] Wu B, Li W, Guidon A, Liu C. Whole brain susceptibility mapping using compressed sensing. Magn Reson Med 2012;67:137–47. doi:10.1002/mrm.23000.

[26] Topfer R, Schweser F, Deistung A, Reichenbach JR, Wilman AH. SHARP edges: Recovering cortical phase contrast through harmonic extension. Magn Reson Med 2015;73:851–6. doi:10.1002/mrm.25148.

[27] Rasmussen KGB, Kristensen MJ, Blendal RG, Ostergaard LR, Plocharski M, O’Brien K, et al. DeepQSM - Using Deep Learning to Solve the Dipole Inversion for MRI Susceptibility Mapping. BioRxiv 2018:278036. doi:10.1101/278036.

[28] Yoon J, Gong E, Chatnuntawech I, Bilgic B, Lee J, Jung W, et al. Quantitative susceptibility mapping using deep neural network: QSMnet. NeuroImage 2018;179:199–206. doi:10.1016/j.neuroimage.2018.06.030.

[29] Liu J, Liu T, de Rochefort L, Ledoux J, Khalidov I, Chen W, et al. Morphology enabled dipole inversion for quantitative susceptibility mapping using structural consistency between the magnitude image and the susceptibility map. NeuroImage 2012;59:2560–8. doi:10.1016/j.neuroimage.2011.08.082.

[30] Ronneberger O, Fischer P, Brox T. U-Net: Convolutional Networks for Biomedical Image Segmentation. Med. Image Comput. Comput.-Assist. Interv. - MICCAI 2015, Springer, Cham; 2015, p. 234–41. doi:10.1007/978-3-319-24574-4_28.

[31] Kingma D, Ba J. Adam: A method for stochastic optimization. ArXiv Prepr ArXiv14126980 2014.

[32] Srivastava N, Hinton GE, Krizhevsky A, Sutskever I, Salakhutdinov R. Dropout: a simple way to prevent neural networks from overfitting. J Mach Learn Res 2014;15:1929–1958.

[33] Jacobsen N, Deistung A, Timmann D, Goericke SL, Reichenbach JR, Güllmar D. Analysis of intensity normalization for optimal segmentation performance of a fully convolutional neural network. Z Für Med Phys 2018. doi:10.1016/j.zemedi.2018.11.004.

[34] Rossum G. Python Reference Manual. Amsterdam, The Netherlands, The Netherlands: CWI (Centre for Mathematics and Computer Science); 1995.

[35] Abadi M, Agarwal A, Barham P, Brevdo E, Chen Z, Citro C, et al. Tensorflow: Large-scale machine learning on heterogeneous distributed systems. ArXiv Prepr ArXiv160304467 2016.

[36] Francois C. Keras 2015. https://github.com/fchollet/keras.

[37] Jones E, Oliphant T, Peterson P. SciPy: Open source scientific tools for Python. 2001.

[38] Oliphant TE. Guide to NumPy. 2nd ed. USA: CreateSpace Independent Publishing Platform; 2015.

[39] Walt S van der, Schönberger JL, Nunez-Iglesias J, Boulogne F, Warner JD, Yager N, et al. scikit-image: image processing in Python. PeerJ 2014;2:e453. doi:10.7717/peerj.453.

[40] Brett M, Hanke M, Markiewicz C, Côté M-A, McCarthy P, Ghosh S, et al. nipy/nibabel: 2.3.0. Zenodo; 2018. doi:10.5281/zenodo.1287921.

[41] Hunter JD. Matplotlib: A 2D Graphics Environment. Comput Sci Eng 2007;9:90–5. doi:10.1109/MCSE.2007.55.

[42] Wang Z, Bovik AC, Sheikh HR, Simoncelli EP. Image Quality Assessment: From Error Visibility to Structural Similarity. IEEE Trans Image Process 2004;13:600–12. doi:10.1109/TIP.2003.819861.

[43] Ravishankar S, Bresler Y. MR Image Reconstruction From Highly Undersampled k-Space Data by Dictionary Learning. IEEE Trans Med Imaging 2011;30:1028–41. doi:10.1109/TMI.2010.2090538.

[44] Langkammer C, Schweser F, Shmueli K, Kames C, Li X, Guo L, et al. Quantitative susceptibility mapping: Report from the 2016 reconstruction challenge. Magn Reson Med 2017:n/a–n/a. doi:10.1002/mrm.26830.

[45] Robinson SD, Dymerska B, Bogner W, Barth M, Zaric O, Goluch S, et al. Combining phase images from array coils using a short echo time reference scan (COMPOSER). Magn Reson Med 2017;77:318–27. doi:10.1002/mrm.26093.

[46] Grodzki DM, Jakob PM, Heismann B. Ultrashort echo time imaging using pointwise encoding time reduction with radial acquisition (PETRA). Magn Reson Med 2012;67:510–8. doi:10.1002/mrm.23017.

[47] Smith SM. Fast robust automated brain extraction. Hum Brain Mapp 2002;17:143–55. doi:10.1002/hbm.10062.

[48] Li W, Avram AV, Wu B, Xiao X, Liu C. Integrated Laplacian-based phase unwrapping and background phase removal for quantitative susceptibility mapping. NMR Biomed 2014;27:219–27. doi:10.1002/nbm.3056.

[49] Liu Z, Kee Y, Zhou D, Wang Y, Spincemaille P. Preconditioned total field inversion (TFI) method for quantitative susceptibility mapping. Magn Reson Med 2016:n/a–n/a. doi:10.1002/mrm.26331.

[50] Sun H, Ma Y, MacDonald ME, Pike GB. Whole head quantitative susceptibility mapping using a least-norm direct dipole inversion method. NeuroImage 2018;179:166–75. doi:10.1016/j.neuroimage.2018.06.036.

[51] Sharma SD, Hernando D, Horng DE, Reeder SB. Quantitative susceptibility mapping in the abdomen as an imaging biomarker of hepatic iron overload. Magn Reson Med 2015;74:673–83. doi:10.1002/mrm.25448.

